# Phylogenomics supports a Cenozoic rediversification of the “living fossil” *Isoetes*

**DOI:** 10.1101/2019.12.23.886994

**Authors:** Daniel Wood, Guillaume Besnard, David J. Beerling, Colin P. Osborne, Pascal-Antoine Christin

**Author notes:** Corresponding author (DW). Molecular Ecology and Fisheries Genetics Laboratory, Bangor University, Deiniol Road, Bangor, Gwynedd, LL57 2UW (Daniel Wood).

## Abstract

The fossil record provides an invaluable insight into the temporal origins of extant lineages of organisms. However, establishing the relationships between fossils and extant lineages can be difficult in groups with low rates of morphological change over time. Molecular dating can potentially circumvent this issue by allowing distant fossils to act as calibration points, but rate variation across large evolutionary scales can bias such analyses. In this study, we apply multiple dating methods to genome-wide datasets to infer the origin of extant species of *Isoetes*, a group of mostly aquatic and semi-aquatic isoetalean lycopsids, which closely resemble fossil forms dating back to the Triassic. Rate variation observed in chloroplast genomes hampers accurate dating, but genome-wide nuclear markers place the origin of extant diversity within this group in the mid-Paleogene, 45-60 million years ago. Our genomic analyses coupled with a careful evaluation of the fossil record indicate that despite resembling forms from the Triassic, extant *Isoetes* species do not represent the remnants of an ancient and widespread group, but instead have spread around the globe in the relatively recent past.

## Introduction

Determining the evolutionary relationships and divergence times between lineages is crucial for understanding the processes that generate diversity and evolutionary novelty [1–3]. Fossils provide a glimpse of the past, by preserving the anatomical features of organisms that existed millions or hundreds of millions of years ago. The fossil record is however very incomplete and often needs to be combined with analyses of extant diversity to infer periods of diversification and extinction of different lineages [4]. A fossil assigned to a lineage of organisms based on shared morphology provides a minimal age for the group, and can therefore help date evolutionary events. Putative causal factors can then be inferred for these events, such as the Chicxulub meteorite impact and the disappearance of the non-avian dinosaurs [5], or the co-incident radiation of angiosperm and insect lineages [6]. In many cases, however, fossils assignable to particular extant lineages of organisms are unavailable, because of a lack of readily fossilisable tissues (e.g. jellyfish) or because the organisms live in environments that do not favour fossilization (e.g. cacti). In addition, morphological traits preserved in fossils may not vary sufficiently to distinguish multiple extant lineages, preventing a precise assignment of the fossils [7–9]. When this pattern of conservation concerns a large number of morphological traits, the extant species are referred to as “living fossils”, a category that includes the coelacanth, cycads, sturgeons, platypus and lungfish that closely resemble fossils from the Mesozoic [10–14]. Clearly a lack of morphological change does not preclude changes in traits poorly represented in the fossil record, such as biochemical or behavioural changes – nevertheless, their unusually conserved morphology through time has long attracted the interest of biologists [15, 16]. This morphological stasis is often associated with decline – with current distributions of “living fossil” taxa interpreted as the remnants of larger ancestral ranges [17, 18], and extant species being the last members of ancient lineages diverging long in the past [19, 20]. However, the morphological uniformity and the resulting difficulties in fossil assignment mean that these hypotheses are difficult to test from the fossil record alone.

Analyses of DNA sequences over the last few decades have resolved the phylogenetic relationships between many extant lineages, and large numbers of selectively neutral changes in the genomes allowed inferring accurate phylogenies even for the most morphologically uniform organisms [21, 22]. Molecular divergence in parts of the tree with informative fossils can then be used to time-calibrate molecular changes in the rest of the tree, allowing inference of divergence times of groups of organisms lacking an appropriate fossil record [23–26]. This molecular dating technique has been used to investigate the evolutionary dynamics of some “living fossil” groups, which in some cases indeed showed ancient within-group divergence and extant distributions resulting from continental drift tens of millions of years ago [17]. In groups such as cycads [13], bryophytes [27] and *Ginkgo* [26], however, extant diversity originated more recently than their conserved morphology would suggest, indicating complex evolutionary dynamics for some “living fossil” taxa. Age estimates from molecular dating techniques remain however sensitive to the treatment of fossil data, variability in the rates of nucleotide substitutions between molecular markers and species, and the correct alignment of nucleotide markers [24, 28–33]. These problems are exacerbated when the only available calibration points are distant from the group of interest, as is by definition the case for “living fossils” [28, 34–37]. Each possible source of error therefore needs to be isolated and carefully considered.

The lycopod genus *Isoetes* exemplifies many of the problems of “living fossil” taxa. The genus has long been of interest due to its status as the last lineage of the isoetalean lycopods. This group, known from at least the late Devonian, dominated terrestrial floras in the Carboniferous [38]. The extant *Isoetes* genus is a small herbaceous aquatic or semi-aquatic plant, generally lacking a stem and consisting of a number of stiff leaves atop a woody corm [39]. It demonstrates a number of unusual features such as roots comparable to fossil stigmarian rootlets [40] and aquatic Crassulacean Acid Metabolism (CAM) [41]. Fossils resembling the *Isoetes* growth form are found in the Triassic onwards, although their exact affinities and relationships to *Isoetes* are unclear [42–44]. A variety of morphological features (such as sunken sporangia, an elaboration of the basal part of the ligule into a glossopodium, and a velum or labium covering the sporangium) that characterise extant *Isoetes* appear at this time, although no single fossil displays all of these features [44]. The appearance of *Isoetites rolandii* in the Jurassic represents the earliest clear example of a isoetalean lycopsid containing all the major features uniting modern *Isoetes*, including the loss of both vegetative leaves and an elongating stem [44, 45], although elongated-stem forms such as *Nathorstiana* persisted until the Early Cretaceous [46]. Fossils of the modern *Isoetes* growth form such as *I. horridus* are subsequently found from the Early Cretaceous and into the Tertiary [42, 44]. The morphology of *Isoetes* appears to have persisted virtually unchanged since at least the Jurassic, and the general growth habit in the lineage is potentially as old as the Triassic.

The close resemblance of fossil taxa to modern *Isoetes* suggests the extant species could be the remnants of a very ancient genus. However, establishing the relationship between these fossils and modern *Isoetes* has proven difficult due to the highly conserved morphology of the genus [18, 38, 44, 47]. The extant 250 *Isoetes* species have a global distribution, yet display very little morphological variation – spore morphology is currently used to distinguish extant species, but reliance on a single character trait makes a classification system vulnerable to homoplasy, convergent evolution and is complicated by widespread hybridisation within the genus [39, 47, 48]. Morphology and the fossil record alone therefore provide limited insights into the relationships among extant and fossil species of *Isoetes,* restricting our ability to understand the temporal origins of extant *Isoetes* species diversity.

The relationships between extant species of *Isoetes* have been inferred using molecular phylogenetics [18, 49, 50], but attempts at linking fossils and extant species have not always been successful. Taylor and Hickey [39] hypothesised based on shared leaf morphology that a small group of South American species and fossil *Isoetes* represented a basal split within the genus, but molecular phylogenetics falsified this hypothesis for extant species [18, 49].

Recent molecular dating studies suggest an origin of extant *Isoetes* species in the Triassic to Jurassic, with species distributions consistent with the breakup of the Gondwana supercontinent [18, 51, 52]. These studies were, however, based on a limited number of markers, mainly from the chloroplast genomes, where high rate variation can make dating estimates strongly dependent on molecular clock model assumptions [24], a potentially significant source of error given the ancient divergence between *Isoetes* and its sister group *Selaginella* [44, 53]. We therefore decided to re-evaluate the divergence times within *Isoetes* using a combination of phylogenomic methods capturing markers spread across the genomes of numerous land plants.

In this study, we generate transcriptomes and genomic datasets for multiple *Isoetes* species and apply multiple molecular dating approaches to estimate the time to the most recent common ancestor of extant *Isoetes* based on nuclear and plastid genomes. Our results shed new light on the age and evolutionary dynamics of this “living fossil” lineage, and show how careful integration of large genomic datasets can help analyses of groups with a poorly informative fossil record.

## Materials and methods

### General approach

In this study, we selected six *Isoetes* for generating genome-wide DNA datasets. These were selected to capture the deeper divergence events within this group based on previous molecular studies [18]. Analyses of nuclear ribosomal DNA available for a large number of *Isoetes* confirmed that the last common ancestor of the selected species likely corresponds to the last common ancestor of extant *Isoetes*, and the low branch length variability throughout the genus suggests the sequenced species represent a good sample of evolutionary rates within the genus (Fig 1). Sparse taxon sampling has been shown to significantly affect estimated dates in some molecular dating studies [54, 55], although not in every case [56, 57]. Rate heterogeneity likely plays an important role in the effect of sparse taxon sampling on the accuracy of molecular dating, with high levels of rate heterogeneity demanding more sampling [55]. The relatively low levels of rate heterogeneity within *Isoetes* (Fig. 1) suggest it is a suitable group to perform molecular dating with a relatively small number of taxa. To further investigate the impact of our sampling scheme, we reanalysed the dataset of Larsén and Rydin [18], which contains 45 *Isoetes* species including all the major clades identified by previous studies of *Isoetes* [50, 58]. This dataset was reanalysed using the same constraints and BEAST settings as previously, but with the 45 *Isoetes* species used reduced to the 6 closest relatives of our chosen species (*I. asiatica, I. coromandelina, I. drummondii, I. echinospora, I. kirkii* and *I. storkii*). This resulted in an estimated crown date of *Isoetes* of 153.4 Ma (53.9-277.3 95% CI), only a 7% increase compared to the full species sampling. This indicates limited taxon sampling should not substantially alter the estimation of the *Isoetes* crown node date.

**Fig 1.**
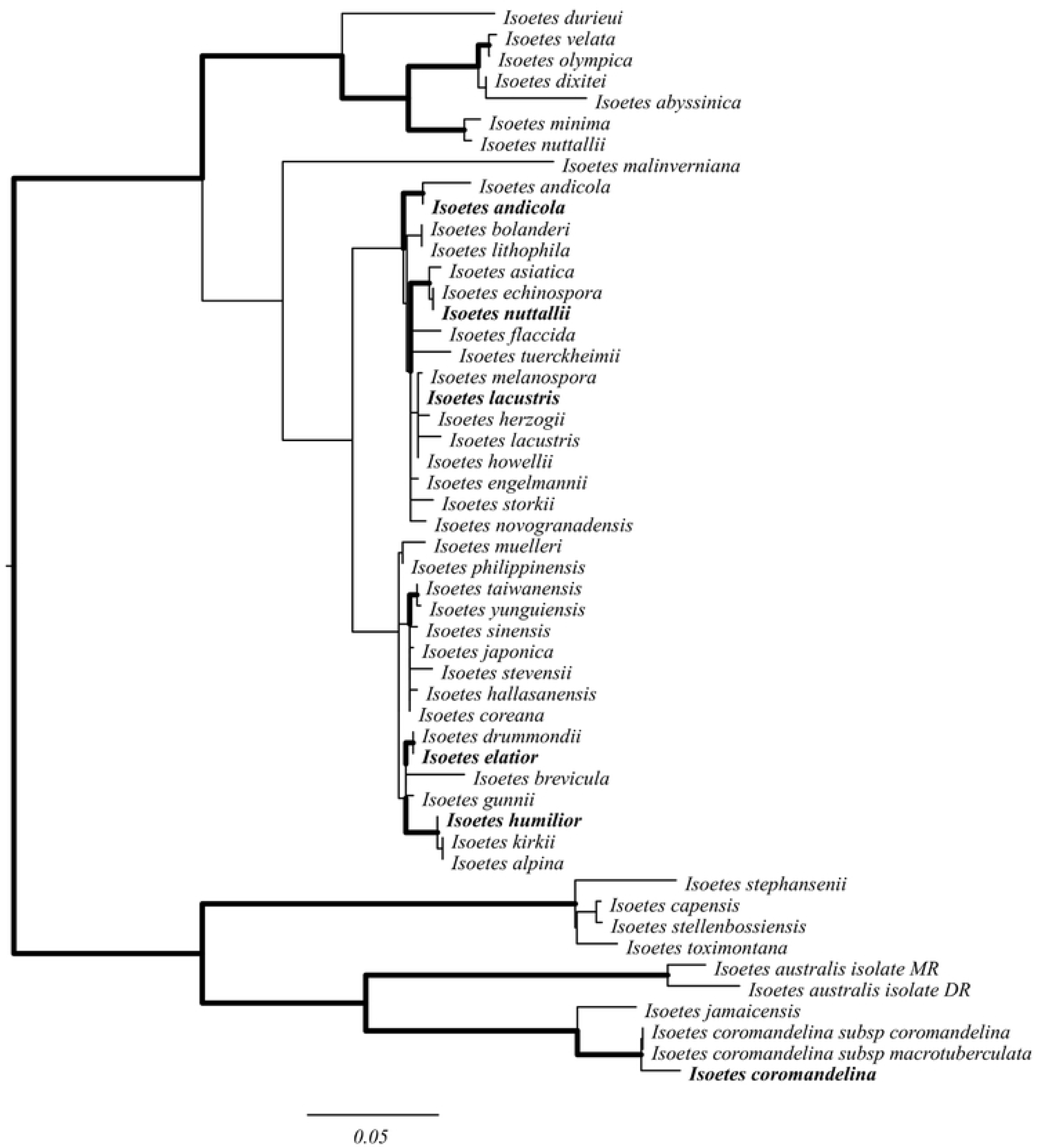
Maximum likelihood phylogram of *Isoetes* nuclear ribosomal internal transcribed spacer. Branch lengths are proportional to the expected number of substitutions per site, with scale bar representing 0.05 substitutions per site. Branches in bold have bootstrap support values greater than 90. Species in bold represent data generated in this study; nuclear ribosomal internal transcribed spacers for other *Isoetes* are from those used in Larsén and Rydin [18].

DNA from these species was then used to generate genome-wide datasets, and different genome partitions were analysed in isolation to get accurate estimates of divergence times. Herbarium specimens represent a useful source of DNA, particularly for globally distributed, hard-to-access groups such as *Isoetes* [59, 60]. Low-coverage whole-genome scans can be applied to these samples, and will yield high coverage for genomic fractions present as multiple copies, such as the organellar genomes [61]. However, highly variable evolutionary rates in chloroplast markers have been reported from seed plants [62, 63], which potentially affect the results of dating methods that differ in their assumptions of rate heterogeneity [24].

Previous studies of the chloroplast marker *rbcL* indicate much higher rates of sequence evolution in *Selaginella* than in *Isoetes* [18, 63, 64], urging for a consideration of nuclear markers, which can be more useful for molecular dating if they show less variation in rates among branches [24]. Genome skimming can provide nuclear sequences, but low coverage makes *de novo* assembly difficult. However, the sequencing reads can be mapped to a reference dataset, providing phylogenetically informative characters [65, 66]. A reference genome is available for *Selaginella*, but it is too distant to allow accurate mapping of reads from *Isoetes*. Transcriptomes provide high coverage of expressed protein-encoding genes, which represent regions of the genome allowing read mapping across distinct species [65, 66]. We consequently decided to generate and assemble a transcriptome for a single *Isoetes* species, which was used as a reference to map reads from low-coverage whole-genome sequencing datasets obtained from the other *Isoetes* species sampled from herbarium collections. The sequencing data were used to obtain chloroplast and nuclear alignments for five *Isoetes* species as well as a number of other land plants (mosses, ferns, lycopods, gymnosperms and angiosperms) sequenced in other studies. The phylogenetic breadth of the datasets allowed the incorporation of fossil evidence providing calibration points spread across the tree.

### Sequence acquisition

Live *Isoetes lacustris* were sampled from Cwm Idwal, Wales and maintained at the University of Sheffield in 40 x 30 x 25 cm transparent plastic containers, with a substrate of sand to a depth of 5 cm, and the containers filled to the top with deionised water. These were placed in a Conviron growth chamber with a 12-h day/night cycle, 495 μmol m^2^s^-1^ light, temperature at 20**°**C during the day and 18**°**C at night, and CO_2_ at 400 ppm for six days. To maximise the number of transcripts retrieved, leaves from three individuals were sampled 3 hours after dark and 3 hours after light and stored immediately in liquid nitrogen. We also generated a transcriptome for *Littorella uniflora* (Plantaginaceae), another species of aquatic plant that shares aquatic CAM photosynthesis [67]. Individuals from this species were also sampled from Cwm Idwal and were grown under a variety of conditions before sampling their leaves as described above. Dried specimens were deposited in the Sheffield University Herbarium (*I. lacustris –* DW1, *L. uniflora –* DW2).

RNA was extracted from the sampled leaves using the RNeasy® Plant Mini Kit (Qiagen), following the manufacturer protocol, with the addition of on-column DNase I digestion (Qiagen RNase-Free DNase Set). We then added 2.5 μl SUPERase-In^TM^ RNase inhibitor (Invitrogen) to 50 μl of extracted RNA to stabilise it. RNA was quantified using a gel electrophoresis, RNA 6000 Nano chips (Aligent) in an Aligent 2100 Bioanalyser, and a Nanodrop 8000. Samples were then prepared for Illumina sequencing using the TruSeq® RNA Sample Prep Kit v2 (Illumina). Paired-end sequencing was performed on an Illumina HiSeq 2500 platform available at Sheffield Diagnostic Genetics Service in rapid mode for 100 cycles, with 24 libraries pooled per lane of flow cell (other samples were from the same or different projects).

DNA from herbarium specimens of five *Isoetes* species were acquired from the DNA Bank from the Royal Botanical Gardens, Kew (Table S2). This was supplemented with one silica gel dried leaf each of *I. lacustris* and *L. uniflora* collected from the field (Cwm Idwal, Wales). Whole genome sequencing of these seven samples was performed at the Genotoul from the University of Toulouse, using previously described protocols [65, 68]. Each sample was sequenced on a 24^th^ of a lane of a flow cell, with other samples from various projects. Raw sequencing reads were cleaned using NGS QC toolkit v2.3.3 [69] by removing adapter sequences, reads with ambiguous bases and reads with less than 80% of positions with a quality score above 20. Low quality bases (q<20) were removed from the 3’ end of remaining reads. Species identity and branch length variability within the genus were assessed by assembling the nuclear ribosomal internal transcribed spacer (nrITS) using NOVOPlasty 2.5.9 [70]. The assembled sequences were aligned to nrITS sequences used in Larsén and Rydin (2015) [18] using MAFFT v7.164 [71]. A phylogeny for this marker was then produced using RAxML v8.2.11 [72], with a GTR + G + I substitution model, identified as the best-fit substitution model through hierarchical likelihood ratio tests.

### Chloroplast data matrix

Cleaned reads from *Isoetes* and *Littorella* corresponding to the chloroplast genomes were assembled using NOVOPlasty, with a 39-bp kmer and a seed sequence of the *I. flaccida* chloroplast genome [64]. In cases where a circular chloroplast genome was not produced, contigs were aligned to the *I. flaccida* chloroplast genome using blastalign [73], and reads corresponding to regions of the reference chloroplast genome not covered by the contigs were used as seed sequences to assemble new contigs. All contigs were subsequently realigned to the reference genome, and overlapping contigs were merged. Chloroplast genome assemblies from 24 additional species representing the major embryophyte taxa, including two *Selaginella* species, were downloaded from NCBI database (Table S3).

Chloroplast protein-coding genes were identified from all chloroplast genomes using DOGMA [74] and coding sequences were extracted using TransDecoder v2.1.0 [75]. A total of 64 genes were identified and aligned by predicted amino acids using t-coffee [76] and MAFFT. Gene alignments were manually inspected and trimmed using AliView [77]. Twelve of them (*clpP, cysaA, psi_psbT, rpl16, rpl21, rps15, ycf1, ycf2, ycf3, ycf10, ycf66, ycf68)* were discarded either due to poor homology or alignment difficulties, or because sequences were obtained from less than 10 of the 33 chloroplast genomes analysed. The remaining 52 chloroplast genes were concatenated, producing a 55,743bp matrix, with 33,496 polymorphic and 25,502 parsimony informative sites. A maximum likelihood phylogeny was generated using RAxML, with a GTR + G + I model of sequence evolution, determined to be the best-fit model using hierarchical likelihood ratio tests. The same matrix was later used for molecular dating.

### Nuclear data matrices

Cleaned RNAseq reads of *I. lacustris* were assembled using Trinity v2.3.2 [75], resulting in 285,613 contigs with an average length of 689 bp. A similar procedure yielded 159,920 contigs for *L. uniflora*, with an average length of 769bp. For each species, the longest open reading frames (ORFs) were extracted using TransDecoder, and for each unigene the contig with the longest ORF was used to build a reference dataset. Cleaned reads from the whole-genome sequencing for each of the *Isoetes* species were then separately mapped to this reference dataset using bowtie2 v2.3.2 [78] in local mode to avoid excluding reads overlapping exon/intron boundaries. Alignments with MAPQ quality below 20 were excluded using SAMtools v1.5 [79]. The SAMtools mpileup utility was then used to generate for each species consensus sequences from the reads mapping to each *I. lacustris* transcript.

Gene duplication and losses are common in nuclear genomes, so a combined reciprocal best blast and phylogenetic approach was adopted to identify groups of co-orthologs covering *I. lacustris* and the other land plants. Families of homologous ORFs generated by the method of Vilella et al. [80] were downloaded from EnsemblPlants. In total, 4,516 homolog families highly conserved among land plants (containing at least one sequence from *Physcomitrella patens, Selaginella moellendorffii, Amborella trichopoda, Oryza sativa, Arabidopsis thaliana* and *Theobroma cacao*) were used for subsequent ortholog identification.

Transcriptome and coding sequence data from seven additional species representing different embryophyte groups were retrieved from the literature [81–87] (Table S4) and ORFs were extracted. Reciprocal best protein BLAST searches assigned ORFs of *I. lacustris, L. uniflora* and the additional embryophyte species to homolog families, with a minimum match length of 50 amino acids and e-value of 10^-7^. The expanded homolog families were then aligned according to their protein sequences using MAFFT, and phylogenies were constructed using RAxML and the GTR + G + I model, which fits most genes and is therefore appropriate for constructing large numbers of gene trees [88–90]. The longest sequence of each monospecific clade of sequences belonging to the same species was kept to remove transcripts representing the same gene or genes that duplicated after the divergence from all other species. The datasets were then realigned and a new phylogeny was inferred. Sets of 1:1 orthologs were then identified as clades containing exactly one gene per species, resulting in 30,258 groups of co-orthologs. Of these, 2,165 contained more than nine species, including *I. lacustris*, *S. moellendorffii* and either *P*. *patens* or *Ceratodon purpurea,* which were needed to use some of the fossil calibration points. These 2,165 orthogroups were realigned, and consensus sequences of the genome skimming data were added to the alignments. Only the 782 orthogroups containing sequences for *I. coromandelina*, which is necessary to capture the earliest split among extant species of *Isoetes* [18] (Fig 1), were considered further. New phylogenetic trees were inferred from these datasets, and genes failing to recover the monophyly of the vascular plants, Isoetopsida (*Isoetes* plus *Selaginella,* 91) or *Isoetes* were considered phylogenetically uninformative and excluded. The remaining 292 datasets were deemed suitable for the phylogenetic problem addressed here, and were used for molecular dating. A phylogenetic tree was inferred separately for each of these markers, and a maximum likelihood phylogeny was also inferred using the 694,437 bp concatenated alignment, which was 41.14 % complete with 443,864 polymorphic and 316,350 parsimony informative sites.

### Calibration points

Time-calibrated trees were inferred from the different markers using the same set of calibration points. To date the crown node of extant *Isoetes*, a fossil constraining the crown node of extant *Isoetes* would be ideal – such a constraint would require a fossil containing a synapomorphy from one of the two descendant branches of this node. Previous studies have identified a geographically diverse group of *Isoetes* including *I. coromandelina* as the outgroup to the rest of the *Isoetes* [18, 50, 58]. Whilst *I. coromandelina* itself contains a number of features initially thought to identify this as a basal member of the group, its presence within “Clade A” identifies these features as derived [18, 49]. No morphological features reliably distinguishing clade A and the rest of the *Isoetes* appear to exist [18, 49].

Therefore, no fossil will contain features distinguishing these two groups, so fossils cannot provide a hard minimum age for this node. Within the *Isoetes* crown group, no morphological features clearly divide different clades [18, 38, 52], therefore using fossil calibrations within the *Isoetes* crown group to produce a hard minimum age is also not possible. The nearest node to the *Isoetes* crown node for which reliable synapomorphies are available is the crown node of the Isoetopsida (*Isoetes* plus *Selaginella*). The Isoeptopsida are a well supported clade appearing in the Devonian, containing synapomorphies such as a heterospory and a ligule (90, 91). Isoetalean lycopsid trees are considered to form a clade within the Isoetales, being more closely related to *Isoetes* than *Selaginella*. This assessment is based on shared synapomorphies including bipolar growth from a shoot like “rhizomorph” structure and secondary woody tissue [44]. Arborescent lycopsids are known from the Frasnian [93, 94] (382.7-372.2 Ma), although the rhizomorph root structure could not be identified in these early fossils. However, discovery of a putatively homosporous arborescent lycopsid (the Isoetales are heterosporous) suggests that arborescence could be a convergent phenotype within the lycopods [95]. As multiple examples of isoetalean arborescent lycopsids, including rhizomorphs, are known from Famennian strata [96, 97] (358.9 to 372.2 Ma), a minimal age of 358 Ma was implemented using a uniform distribution between 358 and 485 Ma.

A maximum age constraint of the crown node of all land plants was set based on the appearance of cryptospores in the fossil record. These abundant spores are considered a likely synapomorphy of early land plants [98]. Their appearance in the fossil record is therefore likely to occur soon after the origins of land plants, making them appropriate for setting a maximum age for land plants [18, 99]. The earliest unequivocal cryptospores are found in the early Middle Ordovician [100] (473-471 Ma). However, pre-Middle Ordovician terrestrial sediments are rare [101], and as no unequivocal cryptospores are found in pre-Ordovician rocks [99, 102], the beginning of the Ordovician (485 Ma) was used as a conservative upper limit for the age of land plants. This maximum age was used to constrain the crown node of the liverworts plus the rest of vascular plants in the chloroplast dataset, and the crown node of the bryophytes plus vascular plants in the nuclear dataset. The minimum age of the same node in both cases was constrained by the early vascular plant macrofossil, *Baragwanathia longifolia* from the Ludlow epoch in the Silurian at 421 Ma [103–105]. This is of a similar age to other putative vascular plant fossils such as *Cooksonia* in the Wenlock epoch [106–108].

For the chloroplast dataset, trees were rooted by constraining each of the liverworts and the rest of the land plants to be monophyletic [104]. For the nuclear dataset, which only contained bryophytes and vascular plants, the tree was rooted by enforcing the monophyly of each of these two groups. Details of fossil calibrations as outlined in Parham et al. [109] are outlined in Table S5.

### Molecular Dating Software

Molecular dating was performed using r8s [110] and BEAST [111], two commonly used relaxed-clock methods that differ in their general approach and the strategy used to assign rates to internal branches of the phylogeny. r8s implements a semiparametric method that uses a penalised likelihood approach to assign rates among branches [112]. The smoothing parameter, which determines the extent to which rates vary among branches, is determined for each dataset using an empirical approach [110]. The method takes a phylogram as input, assumes no uncertainty in topology, and uses a simplified model of nucleotide substitution. BEAST implements a highly parametrised Bayesian method that samples trees generated from nucleotide data using an explicit model of sequence evolution [113]. When using the relaxed molecular clocks implemented in BEAST, rates are uncorrelated across the tree, but an overall distribution of rates is assumed.

For r8s, version 1.81 was used, with the “TN” algorithm and additive penalty function. Cross validation was performed for a range of smoothing parameters from 10^-2^ to 10^6^, increasing by a power of 10^0.5^ each time, and the best smoothing parameter was used for molecular dating. Confidence intervals were obtained by generating 100 bootstrap pseudo-replicates using seqboot [114] and obtaining branch lengths for each of these using RaxML while constraining the trees to the topology generated by the original dataset. These trees were then individually dated using r8s, providing a distribution of ages across the pseudo-replicates. This approach was used to date the chloroplast dataset, the concatenated nuclear dataset, as well as individual nuclear markers.

For BEAST, version v1.8.4 was used. A lognormal relaxed clock was adopted with a GTR + G + I model of nucleotide substitution with four rate categories and a birth-death speciation prior. For the concatenated chloroplast markers, four independent analyses were run for at least 20,000,000 generations and appropriate burn-in periods (at least 10%) were assigned by inspection of the traces using Tracer v1.6 [111]. For individual nuclear genes, BEAST was run for 3,000,000 generations (based on observing convergence times with a subset of genes) with a burn-in period of 50%. Dating the concatenated nuclear dataset was computationally too intensive with this approach. We therefore randomly subsampled 55,743bp (the length of the chloroplast alignment) from the 694,437bp nuclear alignment eight times, and analysed these subsamples with BEAST. The same parameters as the individual nuclear genes were used, with the exception that BEAST was run twice for 10,000,000 generations for each alignment, with a burn-in period of 10%, with convergence verified using Tracer. For comparison we performed r8s on these alignments as described previously.

## Results

### Phylogenetic reconstruction using *Isoetes* nrITS sequences

The maximum likelihood phylogeny based on nrITS broadly agrees with phylogenies previously inferred for the group (Fig 1) [18], and confirms that the species sampled for genomic analyses encapsulate the first divergence within the *Isoetes* species for which molecular data is available (Fig 1). These six samples are moreover representative of the group in terms of branch lengths, and therefore evolutionary rates (Fig 1). The samples sequenced in this study generally cluster with other individuals from the same species sequenced previously, with the exception of *Isoetes nuttallii*. The newly sequenced sample of *I. nuttallii* groups with *I. asiatica* and *I. echinospora,* disagreeing with the topologies found in other studies [18, 49]. The sample in this study was collected in Alaska, USA, where the ranges of *I. nuttallii* and *I. echinospora* overlap [115], making misidentification or a hybrid origin of this individual the likely cause of this discrepancy. We therefore refer to this sample as *I.* sp. throughout the rest of this manuscript and in supplementary data files.

### Phylogenetic reconstruction and dating based on the chloroplast genome

The maximum likelihood phylogeny based on chloroplast markers recapitulated major land plant relationships and expected relationships within the *Isoetes* clade, with *I. coromandelina* being sister to the rest of samples (Fig 2a). The tree was well resolved, with only the *Ceratophyllum*/eudicot split receiving less than 95% bootstrap support. Branch lengths were highly variable, particularly between *Isoetes* and *Selaginella,* with the latter having accumulated approximately 4.5 times more substitutions than *Isoetes* since their most recent common ancestor (Fig 2a).

**Fig 2.**
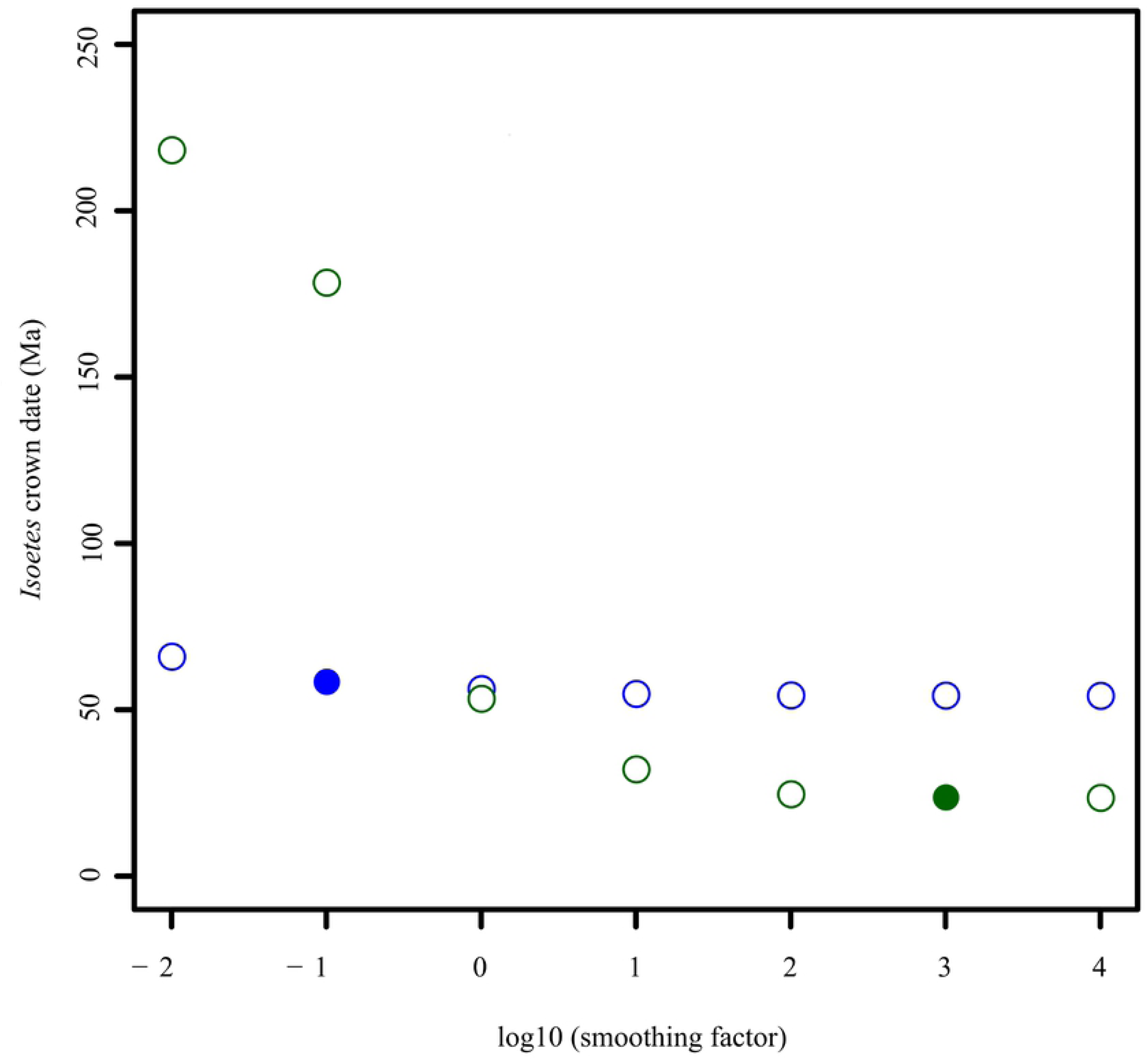
Maximum likelihood phylograms of concatenated chloroplast and nuclear markers. Phylograms are shown for a) concatenated chloroplast markers and b) concatenated nuclear markers. Branch lengths are proportional to the number of expected substitutions per site, with scale bar representing a) 0.07 and b) 0.2 substitutions per site. All bootstraps support values are 100 with the exception of the branch separating *Ananas comosus* from the clade containing *Ceratophylum demersum* in b), which has a support value of 76.

Based on chloroplast markers, r8s estimated the age of the crown group of *Isoetes* at 24.15 Ma with an optimum smoothing parameter of 1000 identified by cross validation, and a 95% bootstrap confidence interval of 23.38-27.40 Ma (Near the Oligocene-Miocene boundary; Table 1). Decreasing the value of the smoothing parameter resulted in an increased age of the *Isoetes* crown group, with a smoothing value of 0.01 giving a crown age of *Isoetes* of 218 Ma (Late Triassic; Fig 3). Whilst low smoothing values result in over-fitted models that perform poorly in cross validation, high levels of smoothing may produce rates that are nevertheless poor predictors of branch lengths in particular parts of the tree. For high smoothing values, the ratio of the effective rate (the branch length divided by the estimated time elapsed) to the rate assigned by the model was 0.33 for the stem branch of *Isoetes* (Fig 4), showing that the branch is significantly shorter than would be expected for the assigned rate and divergence time. On the other hand, the average ratio for the crown branch lengths was 1.33, indicating that the crown branches are longer than would be expected for the assigned rates and divergence times (Fig 4).

**Fig 3.**
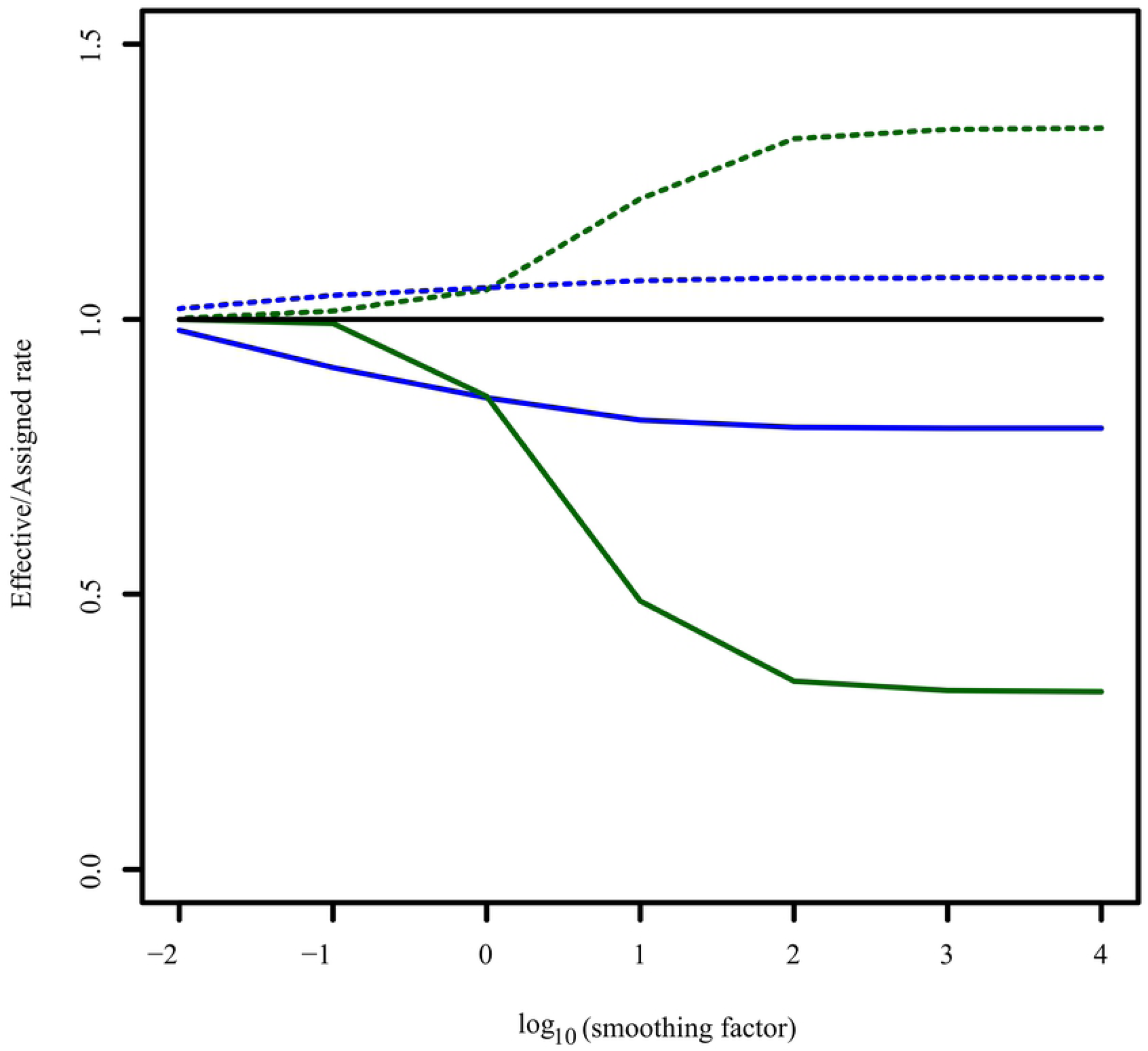
Effect of different smoothing factors on *Isoetes* crown date estimation in r8s. Estimated crown dates for *Isoetes* produced by r8s for concatenated chloroplast (green) and nuclear (blue) datasets are shown for a range of smoothing factors. The best fitting smoothing factor, as identified by cross validation, is highlighted for each dataset by a filled circle.

**Fig 4.**
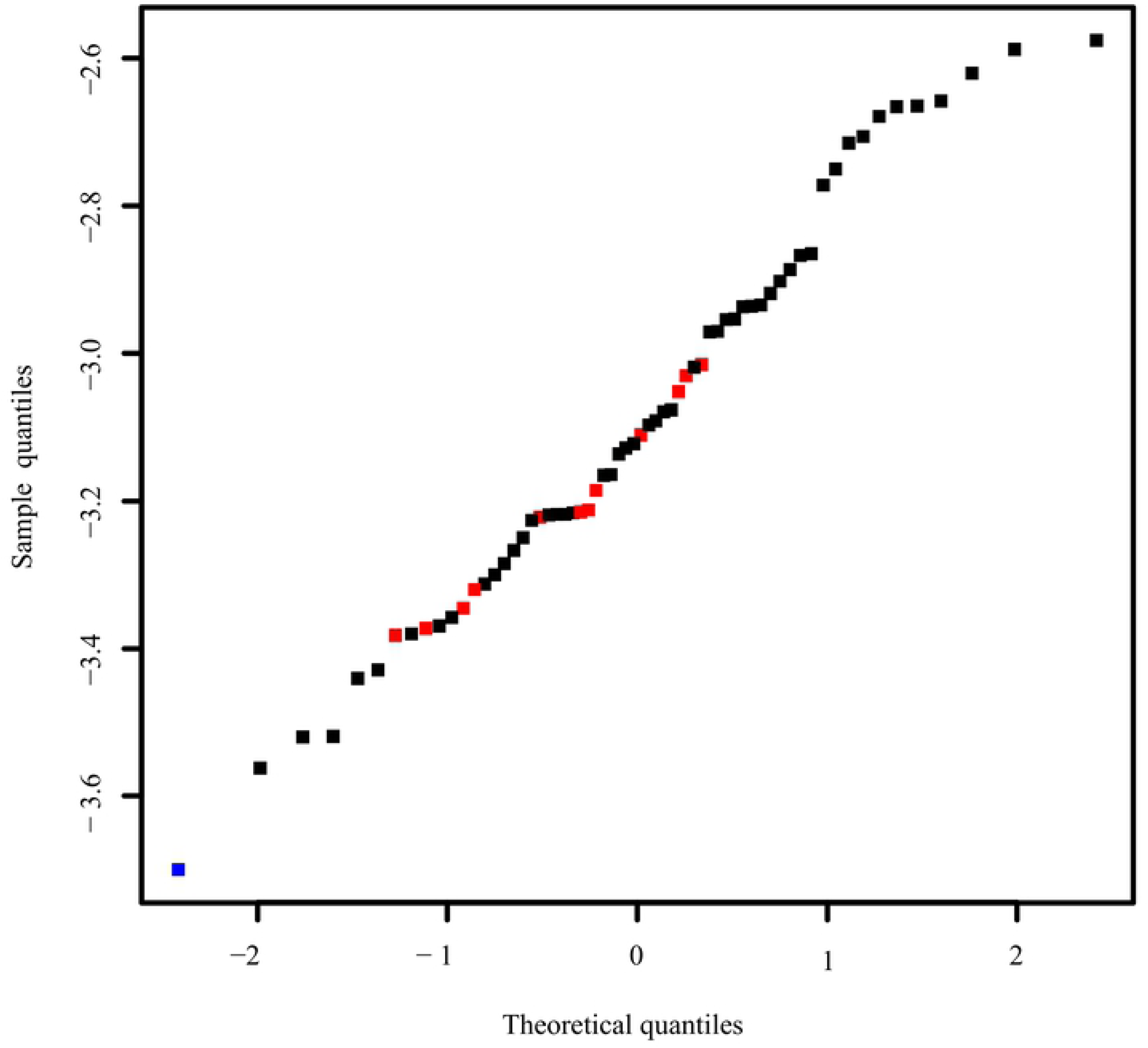
Rate assignment on crown and stem branches of *Isoetes* in r8s. The ratio of effective vs. assigned rates is shown for different smoothing factors in r8s for the stem branch of *Isoetes* (dashed lines) and for the *Isoetes* crown branches (average; solid lines), for the concatenated chloroplast (green) and nuclear (blue) datasets. Solid black line represents effective/assigned rate ratio of 1, for reference.

**Table 1.** Estimates of Isoetes crown date

| Analysis | *Isoetes* crown age - BEAST (95% CI) | *Isoetes* crown age - r8s (95% CI) |
| --- | --- | --- |
| Chloroplast concatenated markers | 28.5 (5.5-51.7) | 24.2 (22.3-25.8) |
| Nuclear concatenated markers | - | 58.9 (57.3-60.1) |
| Nuclear individual markers | 47.6 (24.1, 90.8) | 46.4 (16.1, 85.8) |
| Nuclear concatenated subset | 54.0 (30.3, 82.5) | 62.9 (54.0, 62.9) |

For the same chloroplast markers, BEAST estimated the crown age of *Isoetes* at 28.5 Ma (95% HPD = 5.5-51.7; middle Oligocene; Table 1), similar to the value obtained with the optimum level of smoothing in r8s. Unlike in r8s, rates in BEAST can vary throughout the tree, but their distribution is assigned a priori – in this case a lognormal distribution. Rates in the maximum clade credibility tree accordingly follow a lognormal distribution, with the *Isoetes* stem branch being assigned the lowest rate in the tree and the crown branches assigned rates closer to the average rates in the rest of the tree (Fig 5). For both r8s and BEAST, a date of 23-29 Ma (Oligocene) is obtained via the implicit or explicit inference of a decrease in the rate of evolution along the stem branch, with rates in the crown branches being more similar to those in the rest of the tree. This assumption results from the model, and is not necessarily correct, urging for independent evidence.

**Fig 5.**
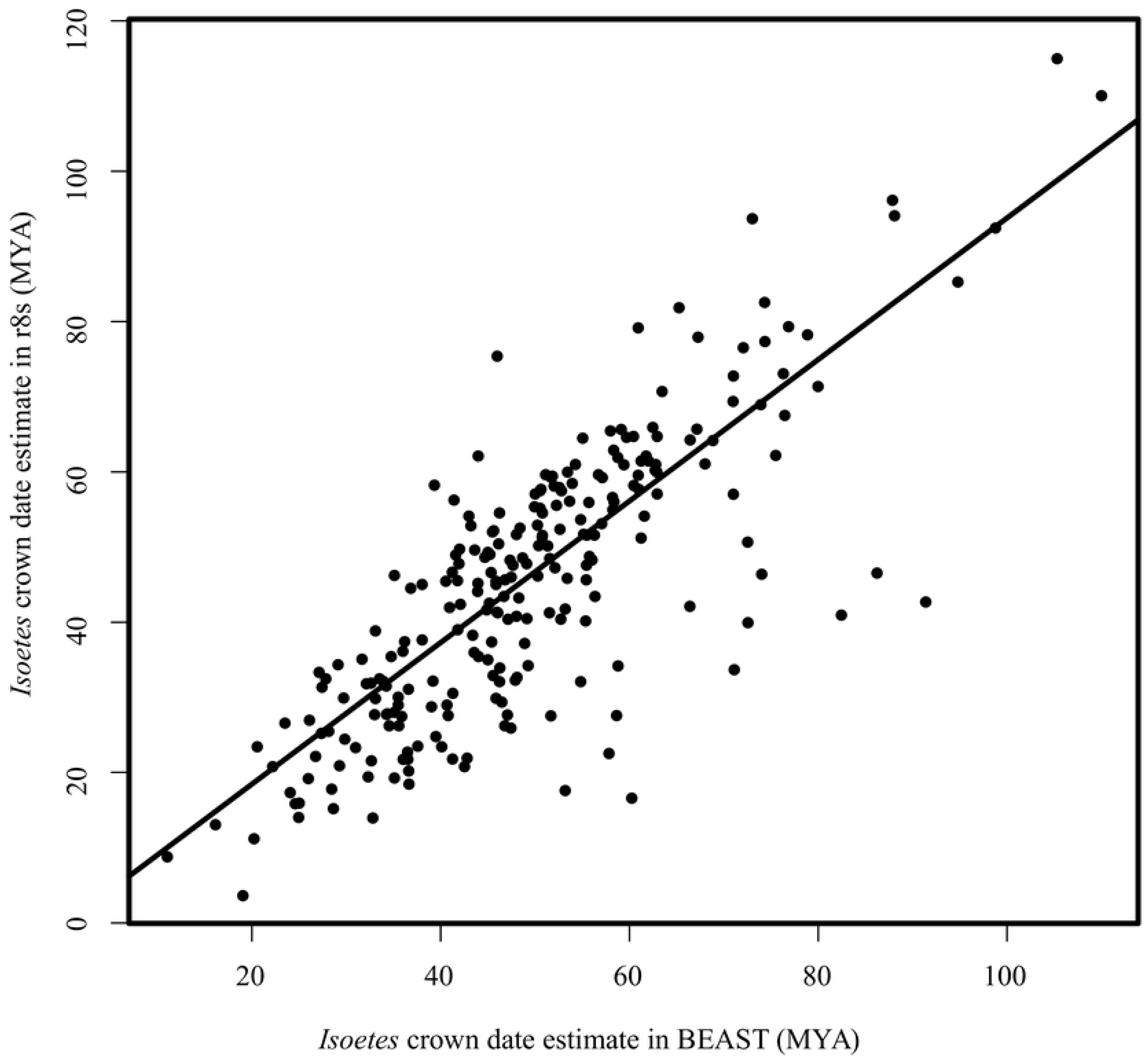
Quantile-quantile plot of BEAST rates for concatenated chloroplast markers. The quantile-quantile plot of log_10_ transformed branch rates is shown for the concatenated chloroplast dataset in BEAST. The values for the Isoetes stem branch (blue) and crown branches (red) are highlighted.

### Phylogenetic reconstruction and dating based on nuclear markers

The concatenated nuclear phylogram also recapitulated major land plant relationships (Fig 2b). The topology of the *Isoetes* clade was consistent with that of the chloroplast phylogeny, with *I. coromandelina* again being sister to all other species. Despite overall longer branch lengths in the concatenated nuclear phylogeny, variation among groups was reduced. Particularly, the total branch lengths from the common ancestor of *Isoetes* and *Selaginella* were much more similar than in the chloroplast phylogeny, with *Selaginella* having accumulated approximately 1.25 times more mutations than *Isoetes* since their last common ancestor (Fig 2b). However, the ratio of the average crown branch length to stem length in the *Isoetes* lineage was very similar between the nuclear and chloroplast markers; approximately 5.8 for the chloroplast dataset and 5.6 for the nuclear dataset (Fig 2).

Dating of the concatenated matrix of nuclear markers in r8s gave an estimated crown node age of *Isoetes* at 58.9 Ma (Table 1), with an estimated stem node age of 358 Ma, at an optimum smoothing value of 0.1. Unlike with the chloroplast markers, the date of the *Isoetes* crown node was similar across all smoothing values tested (Fig 3). Increased smoothing values led to increases in the disparity between effective and assigned rates (Fig. 4), although this was low compared with the concatenated chloroplast alignment (0.82 vs 0.33 for the stem branch and 1.25 vs. 1.33 for the crown branch for a smoothing value of 10^6^). The conservation of the effective rates in the stem and crown branches of *Isoetes* across a range of smoothing parameters indicates that the average rates predicted across the entire nuclear tree are a relatively good fit to both stem and crown branches of *Isoetes* (Fig 4). This suggests that stem and crown branches of *Isoetes* have similar rates, which is consistent with their highly similar length ratios between the chloroplast and nuclear trees (Fig 2).

Dating individual nuclear genes in r8s resulted in a wide range of optimum smoothing values (Fig S1). Low smoothing values frequently resulted in gradient check failures, indicating a single optimum solution is not reached (Fig S1). For genes reaching a single optimum, the median estimated crown date for *Isoetes* was 46.4 Ma with 95% of estimates between 16.1 and 85.8 Ma and 50% of results between 31.9 and 58.3 Ma (Table 1). Overall, the estimated dates form a unimodal distribution (Fig S2). While low values of the smoothing parameter increased the age estimates, all values above 10 yielded estimates centred around 50 Ma, similar to those based on the optimum smoothing values (Fig S2). As with the chloroplast datasets, increasing smoothing values resulted in a decreased effective/assigned stem rate and increased effective/assigned crown rate (Fig S3). The disparities for the optimum smoothing values were again reduced compared to the chloroplast data (Fig S3), indicating the globally optimum smoothing values for the individual nuclear markers fit the stem and crown branches of the *Isoetes* better than in the chloroplast dataset.

Dating individual genes using BEAST gave a median estimate of 47.6 Ma for the crown of *Isoetes*, with 95% of estimates between 24.1 and 90.1 Ma, and 50% between 39.2 and 58.6 Ma. The ages obtained for individual genes were highly correlated between r8s and BEAST (linear model, slope = 0.94, p-value < 0.001; R^2^ = 0.64; Fig 6). Linear modelling suggested a significant but small effect of the percent completeness of the alignments on the estimate for the crown age of *Isoetes*, with a larger effect from the average completeness of *Isoetes* sequences (Table S6). However, the adjusted R^2^ for this latter effect was 0.059 for values obtained with r8s and 0.042 for those obtained with BEAST, indicating that the completeness of the alignment has relatively little impact on the estimated dates. BEAST dating of 55,743bp subsamples of the concatenated dataset gave a mean crown date for *Isoetes* as 54.0 Ma (mean 95% HPD 30.3-82.5Ma; Table 1). The eight individual subsamples gave very similar estimates of the mean crown date, with a standard deviation of 2.8Ma between the different subsamples. r8s gave a slightly older estimate, 62.9 Ma (mean 95% HPD 54.0-62.9; Table 1) with a standard deviation of 2.9Ma between the mean estimates of the different subsamples.

**Fig 6.**
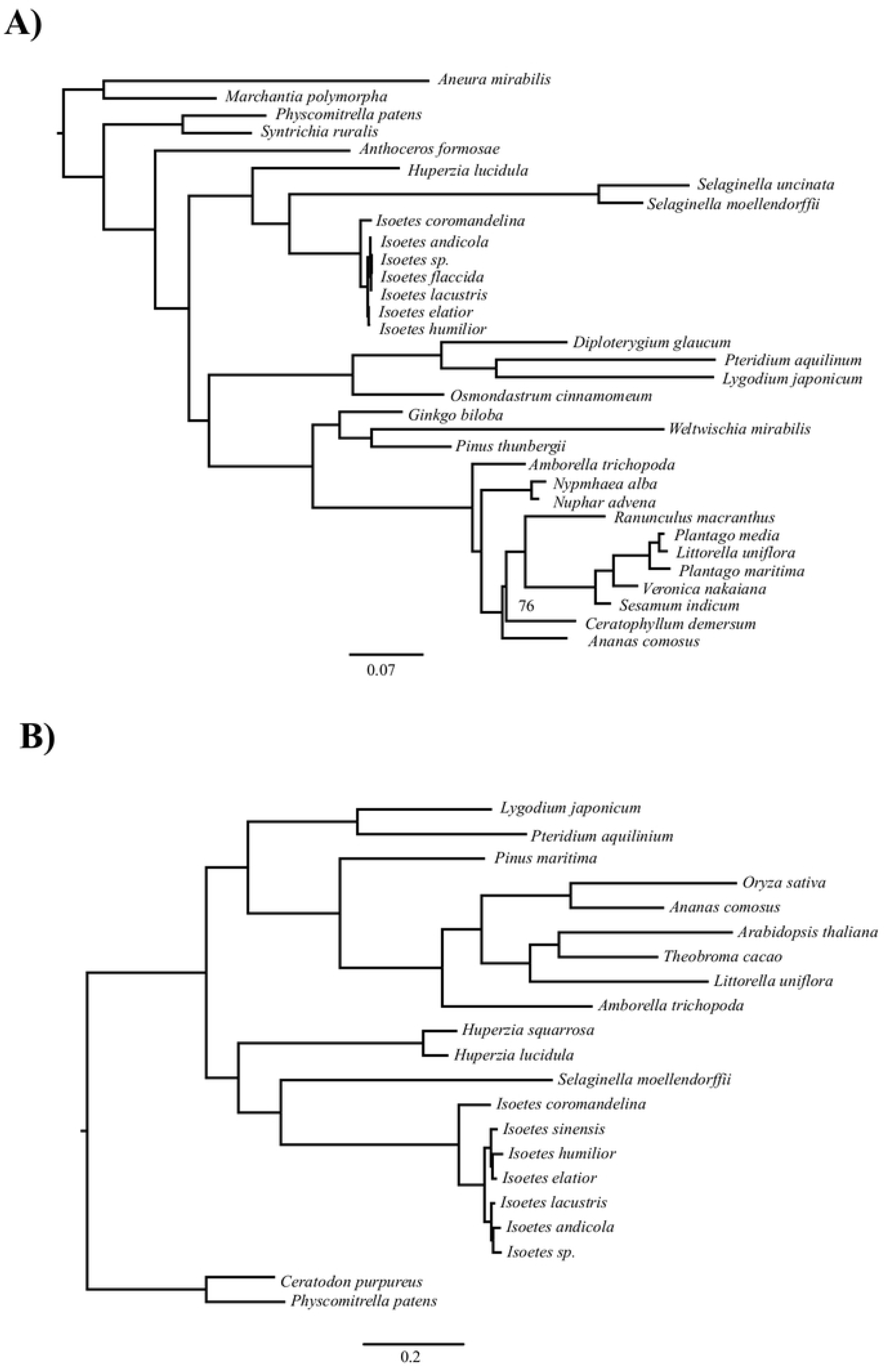
r8s versus BEAST *Isoetes* crown estimates for individual nuclear genes. The scatterplot shows the estimates of the *Isoetes* crown date in r8s and BEAST for each individual nuclear gene. Line represents output of linear model using lm() function in R v3.5.2.

## Discussion

### Nuclear analysis supports a recent origin of extant *Isoetes*

In this study, we used phylogenomics to estimate the age of *Isoetes*, a group of lycopods often interpreted as “living fossils”. Using molecular dating with calibration points on deep branches of the land plant phylogeny, we found very different dates for the crown of *Isoetes* using the chloroplast and nuclear datasets, at 23-24 Ma (Oligocene) and 45-60 Ma (Paleocene and Eocene), respectively (Table 1). These differences are unlikely to be caused by the methods, since BEAST and r8s produced almost identical dates (Table 1; Fig 6), despite the very distinct ways in which these two programs deal with rate variation among branches. Subsets of 55,743bp (the same size as the chloroplast alignment) of the concatenated nuclear alignment gave dates consistent with the other nuclear datasets, indicating alignment size was not the cause of this disparity (Table 1). Instead, the incompatibilities between estimates based on nuclear and chloroplast datasets probably arise from differences in rate variation among branches. Branch lengths varied widely between *Selaginella* and *Isoetes* chloroplast markers (Fig 2a), as previously reported [18, 62–64]. These high levels of variability make low levels of smoothing in r8s relatively poor fits to the data, effectively forcing a uniform rate on the tree that is determined by the average branch length. That, in turn, results in a rate fitted to *Isoetes* that is a poor match to its relatively short branch lengths (Fig 4). As a result, the fitted models predict more changes along the stem branch and fewer changes along the crown branches than occur in the data. Similarly, in BEAST, the lognormal prior distribution results in a relatively low rate assignment on the stem branch compared to the crown branches, which leads to a better fit to the lognormal distribution across branches than if all crown branches had a low rate (Fig 5). We conclude that the high rate variability hampers accurate molecular dating using the chloroplast data. By contrast, the individual and concatenated nuclear datasets have a small disparity between estimated and effective rates, and the crown age estimate for *Isoetes* is consistent across genes (Figs 4 and S2), both in the concatenated versus individual datasets (Table 1) and between BEAST and r8s (Fig 6). The more consistent rates make the nuclear dataset more appropriate for estimating divergence times.

We conclude, based on our nuclear genome-wide analyses, that the diversity of extant *Isoetes* most likely originated during the Paleogene, between 45 and 60 Ma (Paleocene-Ecoene; Table 1), although an origin from the Late Cretaceous to the Micoene is within the 95% confidence interval (16-86Ma; Table 1). This conclusion sharply contrasts with previous estimates of the crown group *Isoetes* of 147-251 Ma (Triassic to Jurassic) [18, 51, 52]. The study of Larsén and Rydin (2015) [18], which found a crown age of 147Ma [96-215 95% CI], was based on three markers, with only *rbcL* aligning with sequences outside of the genus. Removal of the non-coding markers available solely for *Isoetes* species results in a crown estimate for *Isoetes* of 40.5 [22.6-61.6 95% CI] Ma, comparable with the results of the present study (Table S1). Kim and Choi [51] used Triassic *I. beestonii* to provide a narrow lognormal prior with an offset of 245.5, a mean of 1.5 and standard deviation of 0.5Ma for the age of crown *Isoetes,* finding a crown age of 251Ma. Pereira et al. [52] used Jurassic *I. rolandii* to provide a minimum age for the crown node of 145 Ma, finding a crown age of 147Ma (145-154 95% CI). In both these studies, fossils are used to provide strong prior constraints for an ancient date for the crown node of *Isoetes.* However, these fossils do not provide evidence that the split between “clade A” and the rest of *Isoetes* had occurred, as no synapomorphies are known from the extant members of these clades that could be preserved in fossils.

The number of taxa sampled in this study is however lower than in these previous studies. Reduced taxon sampling has been shown to have an impact in some [54, 55], but not all [56, 57], cases, with high levels of rate heterogeneity likely requiring increased taxon sampling [55]. The relatively low levels of rate heterogeneity in *Isoetes* (Fig. 1) indicate this is unlikely to affect our age estimates, and reduction of the taxon sampling in the comprehensive Larsén and Rydin [18] dataset by 87% only resulted in a 7% change in the estimated age of the *Isoetes* crown date (see Materials and Methods). Reanalysis of the entire Larsén and Rydin [18] dataset only with markers alignable outside *Isoetes* resulted in a similar age estimate to the present study, despite the significant differences in taxon sampling (Table S1). These considerations suggest that rather than taxon sampling, the distribution of nucleotide data among groups and the use of fossil evidence are responsible for the differences between this and previous age estimates of *Isoetes*.

Our approach generated nucleotide data that are homogeneously distributed among taxonomic groups, and the fossil evidence is used in a very conservative way, relying solely on external fossils to estimate the age of the crown of *Isoetes*, our group of interest. These considerations allow disentangling the effects of methodological variation, rates of molecular evolution, and treatment of fossils on the molecular dating of a group of “living fossils”.

### Despite morphological stasis, *Isoetes* recently expanded

The relatively young age of the Isoetes crown node indicates that despite morphologically similar forms appearing in the Triassic [42–44], all modern *Isoetes* are descended from a single lineage in the early Cenozoic. This indicates that the fossil *Isoetites* from the Jurassic, and morphologically similar plants from earlier epochs, are likely stem relatives of extant *Isoetes*. The results contrast with the expectation for *“*living fossil” taxa, that extant species members are the last members of once diverse lineages, diverging long in the past [19, 20]. Our results are consistent with an increasing number of studies in groups such as cycads [13], bryophytes [27] and *Ginkgo* [26] showing relatively recent origins of extant species of these groups, despite long periods of morphological stasis.

The global distribution of extant *Isoetes* indicates that this lineage was able to successfully colonise the globe in a relatively short amount of time. This is contrary to the conclusions in previous studies that present *Isoetes* distributions are explained by vicariance due to continental drift [18, 51, 52], based on older estimates for the age of the *Isoetes* crown node. The geographic disparity within several of the major subclades of *Isoetes* - such as Larsen and Rydin’s [18] Clade B containing Mediterranean North American and Indian species – indicates that long distance dispersal events have likely been relatively common in the Cenozoic *Isoetes*. Such a pattern strongly contrasts with the expectation that the distribution of “living fossil” taxa are the remnants of potentially larger ancestral ranges [17, 18]. Our results show that despite having undergone little morphological change for hundreds of millions of years, rather than being the declining remnants of a bygone era, the modern *Isoetes* species instead represent recent arrivals onto the world stage.

## Conclusions

Using molecular dating based on genome-wide datasets and a careful evaluation of the fossil record, we estimated the origins of extant species diversity in *Isoetes*, showing that this group of plants probably diversified in the last 45-60 million years. These results suggest that *Isoetes*-like fossils dating back to the Triassic are stem relatives of extant *Isoetes* species, and that extant *Isoetes* distribution cannot be explained by vicariance from the breakup of Gondwana. Despite their morphological conservatism over hundreds of millions of years, extant *Isoetes* diversified and spread around the world in the relatively recent past, This indicates the morphological stasis of “living fossil” taxa does not preclude lineages of these taxa from diversifying and spreading all around the world

## Acknowledgements

We thank Hélène Holota for help with the sequencing, Luke T. Dunning, Jill K. Olofsson, Jose J. Moreno-Villano and Matheus E. Bianconi for advice in DNA sequencing and computational analyses, Charles Wellman for advice on phylogenetic assignment of cryptospores, and Hannah Sewell for assistance in live plant collection. Catarina Rydin kindly provided the alignment from Larsén and Rydin (2015). PAC is supported by a Royal Society University Research Fellowships (number URF120119).

## Supporting Information

**Figure S1: Optimum Smoothing Values for Nuclear Genes in r8s.** Histogram of optimum smoothing values in r8s identified by cross validation for individual nuclear genes. Proportion of genes for each smoothing value that fail gradient checks are highlighted in red.

**Figure S2: I*s*oetes Crown Dates for Individual Nuclear Genes for different Smoothing Values in r8s.** Histograms showing estimated *Isoetes* crown group dates for individual nuclear genes in r8s that pass gradient checks for a range of assigned smoothing values, and the histogram of estimates where each gene is assigned its optimum smoothing value based on cross validation (final panel).

**Figure S3: Effective/Assigned Rate ratios for Individual Nuclear Genes in r8s.** Histograms of the ratio of effective vs. assigned branch rates for the stem (red) and average value for crown (blue) branches of *Isoetes* for individual nuclear genes in r8s that passed gradient checks for a range of assigned smoothing values, and the histogram of estimates where each gene is assigned its optimum smoothing value based on cross validation (final panel). Median values are displayed in the top righthand corner of each panel.

**Table S1: Reanalysis of dataset of Larsén and Rydin (2015).** The alignment from Larsén and Rydin (2015) was re-analysed using the same constraints and BEAST settings as the previous paper, with at least 3 independent runs reaching ESS > 100.. The dataset contains *rbcL* sequences for *Isoetes* species and other Embryophyte groups, and additional highly variable sequences for *nrITS* and the atpB-rbcL intergenic spacer for *Isoetes* only. *Isoetes* species lacking an rbcL sequence were excluded from the analysis. The entire dataset (i) gave similar estimates of the *Isoetes* crown age to Larsén and Rydin, 2015, but removal of the *atpB-rbcL* intergenic spacer (ii) reduced ages for the Isoetes crown, and removal of either *nrITS* (iii) or both *Isoetes-*specific markers (iv) resulted in ages consistent with the present study.

**Table S2: Kew Herbarium DNA Specimens.** Published with the permission of the Board of Trustees of the Royal Botanic Gardens, Kew.

**Table S3:** Chloroplast Data Sources.

**Table S4:** Transcriptome Data Sources. See main text for references.

**Table S5:** Fossil Constraints Used for Molecular Dating.

**Table S6: Effects of Individual Nuclear Gene Properties on Estimated Dates.** Linear models in R (using the lm function) are used to identify the relationship between a number of alignment properties for individual nuclear genes and the resultant predicted dates in r8s and BEAST. Significant p-values (<0.05) are highlighted in bold.

**FileS1_Individual_Nuclear_Alignments.tar.gz.** Folder containing fasta files for each of the individual nuclear alignments.

**FileS2_Combined_Nuclear_Alignment.tar.gz.** Folder containing fasta file for the combined nuclear alignment.

**FileS3_Chloroplast_Alignment.tar.gz** Folder containing fasta file for the chloroplast alignment.

**FileS4_Chloroplast_Phylogram.tar.gz.** Folder containing nexus file for chloroplast phylogram.

**FileS5_Nuclear_Concatenated_Phylogram.** Folder containing nexus file for concatenated nuclear phylogram.

## References

1. Bromham L, Penny D. The modern molecular clock. Nat. Rev. Genet. 2003;4(3): 216–24.

2. Hipsley CA, Müller J. Beyond fossil calibrations: realities of molecular clock practices in evolutionary biology. Front. Genet. 2014;5: 138.

3. Bromham L, Duchêne S, Hua X, Ritchie AM, Duchêne DA, Ho SYW. Bayesian molecular dating: opening up the black box. Biol. Rev. 2018;93(2): 1165–91.

4. Sauquet H. A practical guide to molecular dating. Comptes Rendus Palevol. 2013;12(6):355–67.

5. Renne PR, Deino AL, Hilgen FJ, Kuiper KF, Mark DF, Mitchell WS et al. Time scales of critical events around the Cretaceous-Paleogene boundary. Science 2013;339(6120): 684–7.

6. Bronstein JL, Alarcón R, Geber M. The evolution of plant–insect mutualisms. New Phytol. 2006;172(3): 412–28.

7. Patterson C. Significance of Fossils in Determining Evolutionary Relationships. Annu. Rev. Ecol. Syst. 1981;12(1): 195–223.

8. Magallón S, Gómez-Acevedo S, Sánchez-Reyes LL, Hernández-Hernández T. A metacalibrated time-tree documents the early rise of flowering plant phylogenetic diversity. New Phytol. 2015;207(2): 437–53.

9. Poinar GO. Chapter Two - The Geological Record of Parasitic Nematode Evolution. In: De Baets K, Littlewood DTJ, editors. Advances in Parasitology. Academic Press, Cambridge, Massachusetts; 2015. pp. 53–92.

10. Smith JL. A living fish of Mesozoic type. Nature 1939;143(3620): 455–6.

11. Hilton EJ, Grande L. Review of the fossil record of sturgeons, family Acipenseridae (Actinopterygii: Acipenseriformes), from North America. J. Paleontol. 2006;80(4): 672–83.

12. Rowe T, Rich TH, Vickers-Rich P, Springer M, Woodburne MO. The oldest platypus and its bearing on divergence timing of the platypus and echidna clades. Proc. Natl. Acad. Sci. 2008;105(4): 1238–42.

13. Nagalingum NS, Marshall CR, Quental TB, Rai HS, Little DP, Mathews S. Recent synchronous radiation of a living fossil. Science 2011;334(6057): 796–9.

14. Kemp A, Cavin L, Guinot G. Evolutionary history of lungfishes with a new phylogeny of post-Devonian genera. Palaeogeogr. Palaeoclimatol. Palaeoecol. 2017;471: 209–19.

15. Darwin C. On the origin of the species by means of natural selection: or, the preservation of favoured races in the struggle for life. London: John Murray; 1859.

16. Lidgard S, Hopkins M. Stasis. Oxford bibliographies on evolutionary biology. New York: Oxford University Press; 2015.

17. Mao K, Milne RI, Zhang L, Peng Y, Liu J, Thomas P et al. Distribution of living Cupressaceae reflects the breakup of Pangea. Proc. Natl. Acad. Sci. USA 2012;109(20): 7793–8.

18. Larsén E, Rydin C. Disentangling the phylogeny of *Isoetes* (Isoetales), using nuclear and plastid data. Int. J. Plant Sci. 2015;177(2): 157–74.

19. Werth AJ, Shear WA. The evolutionary truth about living fossils. Am. Sci. 2014;102(6): 434.

20. Cavin L, Guinot G. Coelacanths as “almost living fossils”. Front. Ecol. Evol. 2014;2: 49.

21. Satler JD, Carstens BC, Hedin M. Multilocus Species delimitation in a complex of morphologically conserved trapdoor spiders (Mygalomorphae, Antrodiaetidae, Aliatypus). Syst. Biol. 2013;62(6): 805–23.

22. Liu Z, Chen G, Zhu T, Zeng Z, Lyu Z, Wang J et al. Prevalence of cryptic species in morphologically uniform taxa – Fast speciation and evolutionary radiation in Asian frogs. Mol. Phylogenet. Evol. 2018;127: 723–31.

23. Arakaki M, Christin P-A, Nyffeler R, Lendel A, Eggli, U, Ogburn RM et al. Contemporaneous and recent radiations of the world’s major succulent plant lineages. Proc. Natl. Acad. Sci. U.S.A. 2011;108(20): 8379–84.

24. Christin P-A, Spriggs E, Osborne CP, Strömberg CAE, Salamin N, Edwards EJ. Molecular dating, evolutionary rates, and the age of the grasses. Syst. Biol. 2014;63(2): 153–65.

25. dos Reis M, Thawornwattana Y, Angelis K, Telford MJ, Donoghue PC, Yang Z. Uncertainty in the timing of origin of animals and the limits of precision in molecular timescales. Curr. Biol. 2015;25(22): 2939–50.

26. Hohmann N, Wolf EM, Rigault P, Zhou W, Kiefer M, Zhao Y et al. *Ginkgo biloba*’s footprint of dynamic Pleistocene history dates back only 390,000 years ago. BMC Genomics 2018;19(1): 299.

27. Laenen B, Shaw B, Schneider H, Goffinet B, Paradis E, Désamoré A et al. Extant diversity of bryophytes emerged from successive post-Mesozoic diversification bursts. Nat. Commun. 2014;5: 5134.

28. Welch JJ, Bromham L. Molecular dating when rates vary. Trends Ecol. Evol. 2005;20(6): 320–7.

29. Rutschmann F, Eriksson T, Salim KA, Conti E, Savolainen V. Assessing calibration uncertainty in molecular dating: The assignment of fossils to alternative calibration points. Syst. Biol. 2007;56(4): 591–608.

30. Goodall-Copestake WP, Harris DJ, Hollingsworth PM. The origin of a mega-diverse genus: dating *Begonia* (Begoniaceae) using alternative datasets, calibrations and relaxed clock methods. Bot. J. Linn. Soc. 2009;159(3):363–80.

31. Ho SY, Phillips MJ. Accounting for calibration uncertainty in phylogenetic estimation of evolutionary divergence times. Syst. Biol. 2009;58(3): 367–80.

32. Brandley MC, Wang Y, Guo X, Montes de Oca AN, Fería-Ortíz M, Hikida T, Ota H. Accommodating heterogenous rates of evolution in molecular divergence dating methods: An example using intercontinental dispersal of *Plestiodon* (Eumeces) lizards. Syst. Biol. 2010;60(1): 3–15.

33. Bromham L. Six Impossible Things before Breakfast: Assumptions, Models, and Belief in Molecular Dating. Trends Ecol. Evol. 2019;34(5):474–86.

34. van Tuinen M, Hedges SB. The effect of external and internal fossil calibrations on the avian evolutionary timescale. J. Paleontol. 2004;78(1): 45–50.

35. Magallón S. Using fossils to break long branches in molecular dating: A comparison of relaxed clocks applied to the origin of angiosperms. Syst. Biol. 2010;59(4): 384–99.

36. Sauquet H, Ho SY, Gandolfo MA, Jordan GJ, Wilf P, Cantrill DJ et al. Testing the impact of calibration on molecular divergence times using a fossil-rich group: the case of *Nothofagus* (Fagales). Syst. Biol. 2011;61(2): 289–313.

37. Duchêne S, Lanfear R, Ho SY. The impact of calibration and clock-model choice on molecular estimates of divergence times. Mol. Phylogenet. Evol. 2014;78: 277–89.

38. Pigg KB. Evolution of Isoetalean lycopsids. Ann. Mo. Bot. Gard. 1992;79: 589–612.

39. Taylor WC, Hickey RJ. Habitat, evolution, and speciation in *Isoetes*. Ann. Mo. Bot. Gard. 1992;79: 613–22.

40. Hetherington AJ, Dolan L. The evolution of lycopsid rooting structures: conservatism and disparity. New Phytol. 2017;215(2): 538–44.

41. Keeley JE. *Isoetes howellii:* A submerged aquatic CAM plant? Am. J. Bot. 1981;68(3): 420–4.

42. Skog JE, Hill CR. The Mesozoic herbaceous lycopsids. Ann. Mo. Bot. Gard. 1992;1:648–75.

43. Retallack GJ. Earliest Triassic origin of *Isoetes* and quillwort evolutionary radiation. J. Paleontol. 1997;71(3): 500–21.

44. Pigg KB. Isoetalean lycopsid evolution: From the Devonian to the present. Am. Fern J. 2001;91(3): 99–114.

45. Ash SR, Pigg, KB. A new Jurassic *Isoetites* (Isoetales) from the Wallowa terrane in Hells Canyon, Oregon and Idaho. Am. J. Bot. 1991;78(12): 1636–42.

46. Karrfalt E. Further observations on *Nathorstiana* (Isoetaceae). American journal of Botany. 1984;71(8): 1023–30.

47. Pfeiffer NE. Monograph of the Isoetaceae. Ann. Mo. Bot. Gard. 1922;9(2): 79–233.

48. Troia A, Pereira JB, Kim C, Taylor WC. The genus *Isoetes* (Isoetaceae): a provisional checklist of the accepted and unresolved taxa. Phytotaxa 2016;277(2): 101–45.

49. Rydin C, Wikström N. Phylogeny of *Isoëtes* (Lycopsida): Resolving basal relationships using *rbcL* sequences. Taxon 2002;51(1): 83–9.

50. Hoot SB, Taylor WC, Napier NS. Phylogeny and biogeography of *Isoëtes* (Isoëtaceae) based on nuclear and chloroplast DNA sequence data. Syst. Bot. 2006;31(3): 449–60.

51. Kim C, Choi HK. Biogeography of North Pacific *Isoëtes* (Isoëtaceae) inferred from nuclear and chloroplast DNA sequence data. J. Plant Biol. 2016;59(4): 386–96.

52. Pereira JBS, Labiak PH, Stützel T, Schulz C. Origin and biogeography of the ancient genus *Isoëtes* with focus on the Neotropics. Bot. J. Linn. Soc. 2017;185(2): 253–71.

53. Banks JA. *Selaginella* and 400 million years of separation. Annu. Rev. Plant Biol. 2009;60: 223–38.

54. Linder HP, Hardy CR, Rutschmann F. Taxon sampling effects in molecular clock dating: an example from the African Restionaceae. Mol. Phylogenet. Evol.. 2005;35(3): 569–82.

55. Soares AE, Schrago CG. The influence of taxon sampling and tree shape on molecular dating: an empirical example from Mammalian mitochondrial genomes. Bioifnorm. Biol. Insights. 2012;6: 129–43.

56. Hug LA, Roger AJ. The impact of fossils and taxon sampling on ancient molecular dating analyses. Mol. Biol. Evol. 2007;24(8): 1889–97.

57. Xiang QY, Thomas DT, Xiang QP. Resolving and dating the phylogeny of Cornales– effects of taxon sampling, data partitions, and fossil calibrations. Mol. Phylogenet. Evol.. 2011;59(1):123–38.

58. Schuettpelz E, Hoot SB. Inferring the root of *Isoëtes*: exploring alternatives in the absence of an acceptable outgroup. Systematic Botany. 2006 Apr 1;31(2): 258–70.

59. Besnard G, Bianconi ME, Hackel J, Manzi S, Vorontsova MS, Christin P-A. Herbarium genomics retrace the origins of C4-specific carbonic anhydrase in Andropogoneae (Poaceae). Bot. Lett. 2018:165(3-4); 419–433.

60. Bieker VC, Martin MD. Implications and future prospects for evolutionary analyses of DNA in historical herbarium collections. Bot. Lett. 2018:165(3-4); 409–18.

61. Bakker, FT, Lei D, Yu J, Mohammadin S, Wei Z, van de Kerke S et al. Herbarium genomics: plastome sequence assembly from a range of herbarium specimens using an Iterative Organelle Genome Assembly pipeline. Biol. J. Linn. Soc. 2016;117(1): 33–43.

62. Bousquet J, Strauss SH, Doerksen AH, Price RA. Extensive variation in evolutionary rate of *rbcL* gene sequences among seed plants. Proc. Natl. Acad. Sci. USA 1992;89(16): 7844–8.

63. Ruhfel BR, Gitzendanner MA, Soltis PS, Soltis DE, Burleigh JG. From algae to angiosperms–inferring the phylogeny of green plants (Viridiplantae) from 360 plastid genomes. BMC Evol. Biol. 2014;14(1): 23.

64. Karol KG, Arumuganathan K, Boore JL, Duffy AM, Everett KD, Hall JD et al. Complete plastome sequences of *Equisetum arvense* and *Isoetes flaccida*: implications for phylogeny and plastid genome evolution of early land plant lineages. BMC Evol. Biol. 2010;10(1): 321.

65. Olofsson JK, Bianconi M, Besnard G, Dunning LT, Lundgren MR, Holota H et al. Genome biogeography reveals the intraspecific spread of adaptive mutations for a complex trait. Mol. Ecol. 2016;25(24): 6107–23.

66. Olofsson JK, Cantera I, Van de Paer C, Hong-Wa C, Zedane L, Dunning LT, Alberti A, Christin PA, Besnard G. Phylogenomics using low-depth whole genome sequencing: A case study with the olive tribe. Mol. Ecol. Resour. 2019;001:1–16.

67. Boston HL, Adams MS. Evidence of crassulacean acid metabolism in two North American isoetids. Aquat. Bot. 1983;15(4):381–386.

68. Lundgren MR, Besnard G, Ripley BS, Lehmann CER, Chatelet DS, Kynast RG et al. Photosynthetic innovation broadens the niche within a single species. Ecol. Lett. 2015;18(10): 021–29.

69. Patel RK, Jain M. NGS QC Toolkit: A toolkit for quality control of next generation sequencing data. PLoS One 2012;7(2): e30619.

70. Dierckxsens N, Mardulyn P, Smits G. NOVOPlasty: de novo assembly of organelle genomes from whole genome data. Nucleic Acids Res. 2017;45(4): e18.

71. Katoh K, Misawa K, Kuma K, Miyata T. MAFFT: a novel method for rapid multiple sequence alignment based on fast Fourier transform. Nucleic Acids Res. 2002;30(14): 3059–66.

72. Stamatakis A. RAxML version 8: a tool for phylogenetic analysis and post-analysis of large phylogenies. Bioinformatics 2014;30(9): 1312–3.

73. Belshaw R, Katzourakis A. *BlastAlign*: a program that uses *blast* to align problematic nucleotide sequences. Bioinformatics 2005;21(1): 122–123.

74. Wyman SK, Jansen RK, Boore JL. Automatic annotation of organellar genomes with DOGMA. Bioinformatics 2004;20(17): 3252–5.

75. Haas BJ, Papanicolaou A, Yassour M, Grabherr M, Blood PD, Bowden J et al. De novo transcript sequence reconstruction from RNA-Seq: reference generation and analysis with Trinity. Nat. Protoc. 2013;8(8): 1494–512.

76. Notredame C, Higgins DG, Heringa J. T-coffee: a novel method for fast and accurate multiple sequence alignment. J. Mol. Biol. 2000;302(1): 205–17.

77. Larsson A. AliView: a fast and lightweight alignment viewer and editor for large datasets. Bioinformatics 2014;30(22): 3276–8.

78. Langmead B, Salzberg SL. Fast gapped-read alignment with Bowtie 2. Nat. Methods 2012;9(4): 357.

79. Li H, Handsaker B, Wysoker A, Fennell T, Ruan J, Homer N et al. The sequence alignment/map format and SAMtools. Bioinformatics 2009;25(16): 2078–9.

80. Vilella AJ, Severin J, Ureta-Vidal A, Heng L, Durbin R, Birney E. EnsemblCompara GeneTrees: Complete, duplication-aware phylogenetic trees in vertebrates. Genome Res. 2009;19(2): 327–35.

81. Der JP, Barker MS, Wickett NJ, dePamphilis CW, Wolf PG. De novo characterization of the gametophyte transcriptome in bracken fern, *Pteridium aquilinum*. BMC Genomics 2011;12(1): 99.

82. Canales J, Bautista R, Label P, Gómez-Maldonado J, Lesur I et al. De novo assembly of maritime pine transcriptome: implications for forest breeding and biotechnology. Plant Biotechnol. J. 2014;12(3): 286–99.

83. Aya K, Kobayashi M, Tanaka J, Ohyanagi H, Suzuki T, Yano K et al. De novo transcriptome assembly of a fern, *Lygodium japonicum*, and a web resource database, Ljtrans DB. Plant Cell Physiol. 2015;56(1): e5.

84. Szövényi P, Perroud PF, Symeonidi A, Stevenson S, Quatrano RS, Rensing SA et al. De novo assembly and comparative analysis of the *Ceratodon purpureus* transcriptome. Mol. Ecol. Resour. 2014;15(1): 203–15.

85. Ming R, Van Buren R, Wai CM, Tang H, Schatz MC, Bowers JE et al. The pineapple genome and the evolution of CAM photosynthesis. Nat. Genet. 2015;47(12): 1435–42.

86. Yang T, Liu X. Comparative transcriptome analysis of *Isoetes sinensis* under terrestrial and submerged conditions. Plant Mol. Biol. Report. 2016;34(1): 136–45.

87. http://www.medplantrnaseq.org/, last accessed 13/11/18

88. Fisher AE, Hasenstab KM, Bell HL, Blaine E, Ingram AL, Columbus JT. Evolutionary history of chloridoid grasses estimated from 122 nuclear loci. Mol. Phylogenet. Evol. 2016;105: 1–14.

89. Dunning LT, Lundgren MR, Moreno-Villena JJ, Namaganda M, Edwards EJ, Nosil P et al. Introgression and repeated co-option facilitated the recurrent emergence of C_4_ photosynthesis among close relatives. Evolution 2017;71(6): 1541–55.

90. Abadi S, Azouri D, Pupko T, Mayrose I. Model selection may not be a mandatory step for phylogeny reconstruction. Nat. Commun. 2019;10(1): 934.

91. Rolle, F. Kryptogamen. In: Förster W, Kenngott A, Ladenburg D, Reichenow A, Shenk D, Schömilch D et al. (eds.). Encyklopaedie der Naturwissenschaften. 1885;12. pp. 211–277.

92. Hao S, Xue J, Wang Q, Liu Z. *Yuguangia ordinata gen. et sp. nov*., a new lycopsid from the Middle Devonian (late Givetian) of Yunnan, China, and its phylogenetic implications. Int. J. Plant Sci. 2007;168(8): 1161–75.

93. Stein WE, Berry CM, Hernick LV, Mannolini F. Surprisingly complex community discovered in the mid-Devonian fossil forest at Gilboa. Nature 2012;483(7387): 78–81.

94. Berry CM, Marshall JEA. Lycopsid forests in the early Late Devonian paleoequatorial zone of Svalbard. Geology, 2015;43(12): 1043–6.

95. Xu HH, Wang Y, Wang Q. A new homosporous, arborescent lycopsid from the Middle Devonian of Xinjiang, Northwest China. Palaeontology 2012;55(5): 957–66.

96. Wang Y, Berry CM. A novel lycopsid from the Upper Devonian of Jiangsu, China. Palaeontology 2003;46(6): 1297–311.

97. Cressler WL, Pfefferkorn HW. A Late Devonian isoetalean lycopsid, *Otzinachsonia beerboweri*, gen. et sp. nov., from north-central Pennsylvania, USA. Am. J. Bot. 2005;92(7): 1131–40.

98. Wellman CH. The invasion of the land by plants: when and where? New Phytol. 2010;188(2), 306–9.

99. Morris JL, Puttick MN, Clark JW, Edwards D, Kenrick P, Pressel S et al. The timescale of early land plant evolution. Proc. Natl. Acad. Sci. USA 2018;115(1): E2274–83.

100. Rubinstein CV, Gerrienne P, de la Puente GS, Astini RA, Steemans P. Early Middle Ordovician evidence for land plants in Argentina (Eastern Gondwana). New Phytol. 2010;188(2): 365–9.

101. Wellman CH, Strother PK. The terrestrial biota prior to the origin of land plants (embryophytes): a review of the evidence. Palaeontology 2015;58(4): 601–27.

102. Kenrick P. Palaeobotany: fishing for the first plants. Nature 2003;425(6955): 248.

103. Garratt MJ, Tims JD, Rickards RB, Chambers TC, Douglas JG. The appearance of *Baragwanathia* (Lycophytina) in the Silurian. Bot. J. Linn. Soc. 1984;89(4): 355–8.

104. Kenrick P, Crane PR. The origin and early evolution of plants on land. Nature 1997;389(6646): 33.

105. Magallón S, Hilu KW, Quandt D. Land plant evolutionary timeline: gene effects are secondary to fossil constraints in relaxed clock estimation of age and substitution rates. Am. J. Bot. 2013;100(3): 556–73.

106. Edwards, D, Feehan, J. Records of *Cooksonia*-type sporangia from Late Wenlock Strata in Ireland. Nature. 1980;287(5777). 41–2.

107. Edwards D, Feehan J, Smith DG. A late Wenlock flora from Co Tipperary, Ireland. Bot. J. Linn. Soc. 1983;86(1-2), 19–36.

108. Libertín M, Kvaček J, Bek J, Žárský V, Štorch P. Sporophytes of polysporangiate land plants from the early Silurian period may have been photosynthetically autonomous. Nature plants. 2018;4(5): 269

109. Parham JF, Donoghue PC, Bell CJ, Calway TD, Head JJ, Holroyd P et al. Best practices for justifying fossil calibrations. Systematic Biology. 2011;61(2): 346–59.

110. Sanderson MJ. r8s: inferring absolute rates of molecular evolution and divergence times in the absence of a molecular clock. Bioinformatics 2003;19(2): 301–2.

111. Drummond AJ, Rambaut A. BEAST: Bayesian evolutionary analysis by sampling trees. BMC Evol. Biol. 2007;7(1): 214.

112. Sanderson MJ. Estimating absolute rates of molecular evolution and divergence times: A penalized likelihood approach. Mol. Biol. Evol. 2002;19(1): 101–9.

113. Drummond AJ, Ho SYW, Phillips MJ, Rambaut A. Relaxed phylogenetics and dating with confidence. PLoS Biol. 2006;4(5): e88.

114. Felsenstein J. {PHYLIP} (Phylogeny Inference Package) version 3.6a3. 2002. Distributed by the author.

115. Britton DM, Brunton DF, Talbot SS. *Isoetes* in Alaska and the Aleutians. Am. Fern J. 1999: 133–41.

